# Altered neuromechanical strategies of the paretic hip and knee joints during a step-up task

**DOI:** 10.1101/2020.12.02.407122

**Authors:** Vatsala Goyal, Andrew Dragunas, Robert L. Askew, Theresa Sukal-Moulton, Roberto López-Rosado

## Abstract

**Background:** Stroke often leads to chronic, neural-derived motor impairments in the paretic lower limb, such as weakness, abnormal extensor torque coupling, and reduced ranges of motion. These impairments can constrain lower extremity movement and negatively impact the ability to navigate uneven terrain. Quantification of biomechanical strategies used by individuals with chronic stroke to step up would offer insight into the neural consequences of a stroke.

**Research Question:** What are the altered kinetic and kinematic strategies of the leading paretic hip and knee joints while swinging and pulling-up onto a step?

**Methods:** A total of 10 participants were included in this mixed design study: 5 adults with hemiparetic stroke and 5 age-matched adults without stroke. Participants were instructed to step up onto a 4-inch platform, where joint kinetic and kinematics of the hip in the frontal plane and the hip and knee in the sagittal plane were quantified. A mixed effects linear regression model with two fixed effects of group (stroke and control) and lower limb (LL: dominant/non-paretic and non-dominant/paretic) was used to compare peak joint torques and angles. Another mixed effects model with two fixed effects of peak hip and knee extension torque was used to investigate whether these main effects could predict peak hip abduction torque.

**Results:** Altered biomechanical strategies of the paretic limb for step ascent included reduced sagittal plane flexion angles during swing, reduced hip abduction and knee extension torque combined with increased hip extension torque during pull-up stance, and abnormal torque coupling between the hip adductors and sagittal plane extensors.

**Significance:** These differences can be linked to the neural consequences of a hemiparetic stroke, including corticospinal damage and upregulation of bulbospinal pathways as compensation. Overall, our findings can inform interventions for individuals with chronic stroke in navigating uneven terrain to maximize daily community activity.

## Introduction

Stroke is the leading cause of disability in the United States [1] and often leads to chronic, asymmetrical lower limb motor impairments that severely limit independent community ambulation. Neural-derived biomechanical deficits include hip and knee weakness, abnormal extensor coupling between the hip adductors and sagittal plane extensors, and reduced ranges of motion in the paretic (more affected) lower limb [2-5]. These impairments can constrain lower extremity movement to maladaptive strategies that are mechanically inefficient. Indeed, individuals with stroke increasingly depend on the non-paretic (less affected) limb for balance and stance loading during steady-state walking [6-8]. However, steady-state walking only constitutes a small part of community ambulation. Maximizing daily occupational, social, and physical activities requires negotiating uneven terrain such as stairs and curbs [9,10].

Compared to level ground walking, stepping up onto a stair or curb requires generation of substantial hip and knee flexion angles during swing to achieve proper toe clearance and greater knee extension moments during the pull-up phase of stance to elevate the body’s center of mass [11-14]. For older adults without a stroke, researchers investigating stair ascent have reported similar peak hip and knee extension torques but greater peak hip abduction torques during pull-up compared to their younger peers [15]. This increase in frontal plane hip torque could be a strategy for older adults to maximize stability and mitigate risk of falling [15]. However, neural-derived weakness in the hip abductors and extensor/adductor torque coupling in the paretic lower limb following stroke may challenge use of this strategy for stability [2-4]. Novak et al. reported similar kinematic profiles but decreases in peak hip abduction and knee extension moments during pull-up in the paretic limb compared to non-paretic and control limbs [16]. Nevertheless, this analysis did not consider the substantial hip and knee flexion angles required to swing the limb onto the step or the extensor/adductor torque coupling that could explain decreases in hip abduction torque generation. Additionally, the authors focused on biomechanics of the second step, whereas the higher hip abduction moments required to transition from standing or level walking to stair ascent [17] make the first step likely more difficult post-stroke. These considerations may reveal the limits of lower extremity neuromuscular control.

The purpose of this study was to quantify the kinetics and kinematics of the leading hip and knee during a step-up task in individuals with and without stroke. We focused on frontal plane metrics in the hip and sagittal plane metrics in the hip and knee to characterize the joints with the greatest contributions to swinging and pulling up onto a step [11-15]. Given joint weakness and reduced ranges of motion in the paretic lower limb, our main hypotheses were: 1) the paretic limb would have decreased peak hip abduction, hip flexion, and knee flexion angles while leading in swing phase compared to the non-paretic and control limbs and 2) the paretic limb would have decreased peak hip abduction, hip extension, and knee extension torque while leading in the pull-up stance phase compared to the non-paretic and control limbs. We also hypothesized that a decrease in peak hip abduction torque during pull-up could be predicted from increases in peak hip extension and knee extension torques in the leading paretic limb, suggesting abnormal extensor coupling. Quantification of these peripheral metrics can offer insight into the permanent neural consequences of a stroke and allow for more targeted interventions to maximize activity in the community.

## Methods

### I. Participant summary

A total of 10 participants were included in this study: 5 adults with hemiparetic stroke (4 with right hemiparesis, 2 females, age = 59.0 ± 7.6 [mean ± SD]) and 5 age-matched adults without stroke (3 females, age = 60.4 ± 5.0). Adults aged 21-65 years who could independently walk up to 1000 ft with a speed of at least 240 ft/min and step up and down a 4-inch platform were eligible for this study. Furthermore, participants with stroke were included if they had a clinical presentation of hemiparesis from a diagnosed unilateral stroke at least one year prior to study participation. Exclusion criteria were 1) orthopedic surgeries or fractures in the trunk, pelvis, or lower limb joints, 2) current use of botulinum toxin injections or other muscle relaxants in the lower extremity, 3) inability to stand or take a step up independently, and 4) health risks from comorbidities. This study was approved by Northwestern University’s Institutional Review Board and all participants provided informed consent.

### II. Experimental Set-Up

Participants performed a series of stair-stepping trials in a gait laboratory. We built two custom-machined 4-inch wooden steps that were placed on individual force plates. These steps were designed to cover the entire force plate surface area so that participants’ center of pressure on the step was appropriately translated to the force plate. A 4-inch step height was selected as it proved sufficiently challenging and achievable for all of our participants with stroke.

A 12-camera motion capture system (Qualisys, Gothenburg, Sweden) and four in-ground force plates (AMTI, Glenview, IL) were used to record the 3D positions of retro-reflective markers and ground reaction forces, respectively. Marker data was recorded at 60 Hz and ground reaction force data was recorded at 1200 Hz. Markers were placed bilaterally on anatomical landmarks of the pelvis, thigh, shank, and foot, including: anterior superior iliac spines, posterior superior iliac spines, greater trochanters, lateral femoral epicondyles, lateral malleoli, calcanei, and the 2nd and 5th metatarsals. A single marker was placed on the sternum and 4-marker tracking clusters were affixed bilaterally to the thigh and shank.

### III. Experimental Procedure

Prior to the stair-stepping trials, a licensed physical therapist administered clinical assessments to evaluate walking and balance function. Clinical measures included the Lower Extremity (LE) Fugl-Meyer [18], the Activities-specific Balance Confidence (ABC) Scale [19], and the Step Test [20]. Lower limb dominance was self-reported for the control group and established as the non-paretic side for the stroke group. Vital signs were taken at prescribed intervals (pre-, during, post-experiment) to ensure participants had an appropriate response to activity.

To minimize the risk of falling, participants were secured with a passive harness (Aretech, Ashburn, VA) attached to an overhead trolley system that allowed for unrestricted anteroposterior movement. Participants with stroke were not allowed to use assistive devices such as canes or walkers during the experiment but were allowed use of their ankle foot orthoses if typically worn for community ambulation. Participants started each trial with their feet on two individual force plates that were adjacent to the force plates with the wooden steps. Outlines of each foot were taped onto these force plates to indicate starting foot position. Prior to data collection, participants first practiced stepping up onto the wooden steps, alternating between leading with their dominant/non-paretic and non-dominant/paretic lower limb. After becoming comfortable with the task, participants were instructed to take a full step up as they normally would while avoiding the use of their hands for assistance. Participants were given breaks when requested and after 5 trials had been completed. Participants completed 12 trials for each lower limb for a total of 24 trials per participant.

### IV. Data Processing & Analysis

Kinematic data were visually inspected in Qualisys Track Manager to confirm that markers were labeled appropriately. Data were then processed in Visual 3D (C-Motion, Germantown, MD) to calculate joint kinetics and kinematics. To account for the step height during inverse dynamic calculations, two virtual force platforms were created in the software, with the 3D position of these platforms defined by 4 markers placed on the corners of each step. Marker and force data were filtered with a 4th order Butterworth filter with a cut-off of 6Hz to remove high-frequency fluctuations in the data. Stepping events for toe-off and heel-strike were determined by a 5 N threshold on each force plate, and these events were then visually inspected to confirm accuracy. We then used inverse dynamics to calculate the following joint torques and angles: hip abduction/adduction, hip flexion/extension, and knee flexion/extension.

All data were exported into MATLAB (MathWorks, Inc., Natick, MA) for further processing. Kinematic data were extracted when the leading lower limb was swinging onto the step, between leading foot toe-off and heel-strike. This window was normalized to 100% of the swing phase and peak angles in hip abduction (HABDa), hip flexion (HFLEX), and knee flexion (KFLEX) were considered for statistical analysis. Kinetic data were normalized to body weight and extracted when the leading lower limb was in stance pulling the body up onto the step, between leading foot heel-strike and trailing foot heel-strike. This window was normalized to 100% of the stance phase and peak torques in hip abduction (HABD), hip extension (HEXT), and knee extension (KEXT) were considered for statistical analysis.

### V. Statistical Analysis

All statistical analysis was done in Stata IC 14.1 (StataCorp LLC, College Station, TX, USA). Independent two-sample t-tests were used to compare height, weight, and ABC scores between the two groups. A paired t-test was used for the Step Test to compare differences in step count between the two lower limbs within each group. Q-Q plots were visually inspected to assess normality of the kinematic and kinetic outcome variables. This study had one between subjects factor of group (stroke and control) and one within subjects factor of lower limb (dominant/non-paretic and non-dominant/paretic) representing a mixed study design. Thus, we ran a mixed effects linear regression model for each outcome variable with two fixed effects of group and lower limb (LL) and one random effect of subject to control for repeated measures within participants. We also ran another mixed effects linear regression model to understand whether the fixed effects of peak HEXT and KEXT torque efforts could predict peak HABD torque generation, with one random effect of subject. The threshold for statistical significance was set a priori to 0.05.

## Results

### I. Participant Metrics

All participant metrics are displayed in Table 1. There was no significant difference between weight (p = 0.846), height (p = 0.335), or ABC score (p = 0.092) between the groups. For the Step Test, the non-paretic lower limb had a significantly greater step count than the paretic lower limb in the stroke group (p = 0.002), but there were no significant differences between the lower limbs for the control group (p = 0.681). The average LE Fugl-Meyer score for the stroke group was just under 21 points (Table 1), indicating a moderate-to-high level of functional mobility [21].

**Table 1.** Mean (SD) participant-specific metrics and clinical assessment outcomes.

|  |  |  |  |  | Step Test |  |
| --- | --- | --- | --- | --- | --- | --- |
|  | Weight<br>(kg) | Height<br>(in) | ABC<br>Score | LE Fugl-Meyer | Dominant/<br>Non-paretic | Non-Dominant/Paretic |
| **Stroke** | 68.6<br>(12.2) | 66.7<br>(3.5) | 84.3<br>(13.9) | 20.4 (5.0) | 11.5 (4.7) | 8.1 (4.1) |
| **Control** | 66.9<br>(12.9) | 64.7<br>(2.6) | 98.0<br>(3.3) | -- | 18.3 (3.7) | 18.0 (3.7) |

### II. Joint Kinetics

Joint torque means and standard deviations are displayed in Table 2 and Figure 1. For HABD, there was no significant main effect of group (p = 0.480) but there was a significant main effect of lower limb (LL) (p = 0.009), where the dominant/non-paretic limbs generated higher peak HABD torques than the non-dominant/paretic limbs. The interaction between group and LL was not significant for HABD (p = 0.594). For HEXT, there was a significant effect of group (p < 0.001), where the stroke group generated higher peak HEXT torques than the control group, but there was not a significant effect of LL (p = 0.480). The interaction term was not significant for HEXT (p = 0.573). For KEXT, there was a significant effect of group (p = 0.031) and LL (p < 0.001). Their interaction was also significant (p < 0.001). The stroke paretic limb exhibited lower peak KEXT torques than the stroke non-paretic limb (Confidence Interval (CI): [0.667 0.994]), control dominant limb (CI: [0.031 0.372]), and control non-dominant limb (CI: [0.009 0.480]) Additionally, the stroke non-paretic limb exhibited greater peak KEXT torques than the control dominant (CI: [−0.850 −0.409]) and control non-dominant limbs (CI: [−0.860 −0.313]).

**Figure 1.**
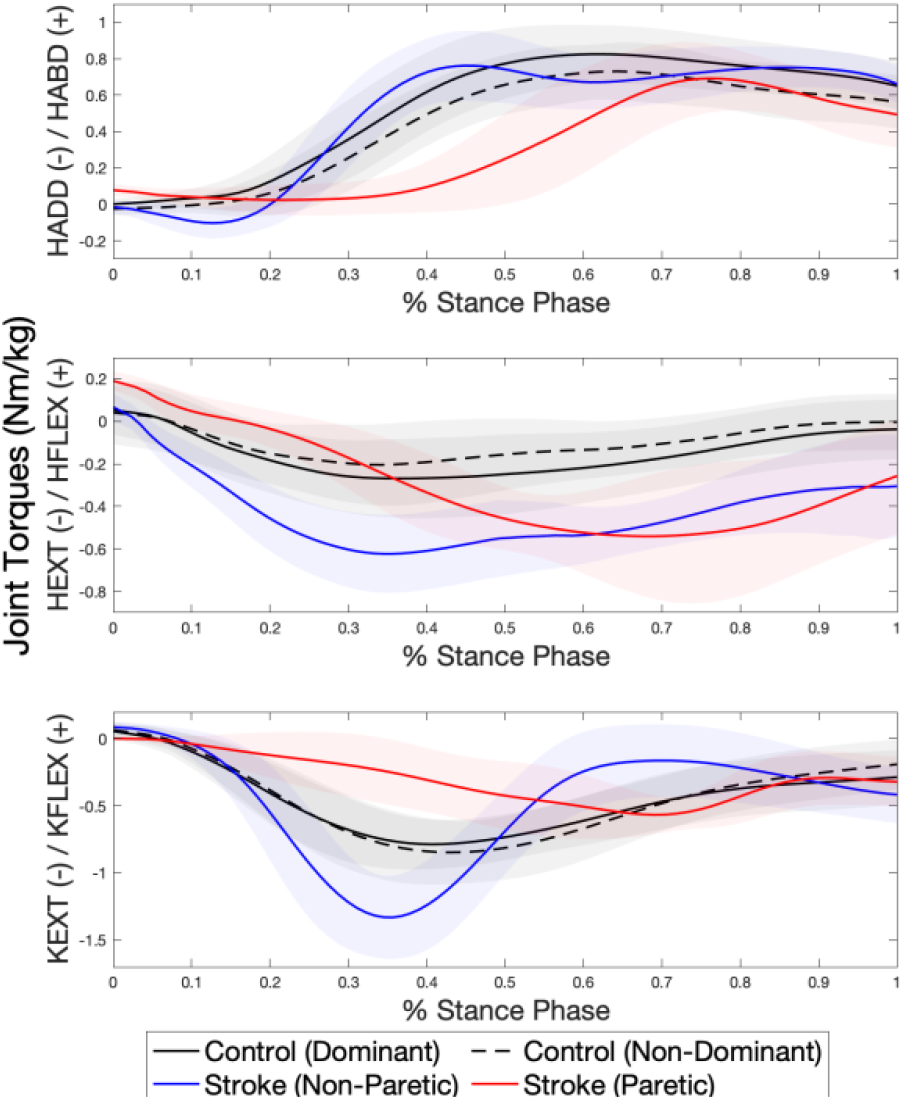
Joint torque traces for the stance phase pulling the body up onto the step. The x-axis of each subplot is time normalized to 100% of the stance phase and the y-axis is joint torque normalized to body weight. Shaded regions represent one standard deviation. Red line = stroke paretic limb, blue line = stroke non-paretic limb, black dashed line = control non-dominant limb, black solid line = control dominant limb.

**Table 2.**
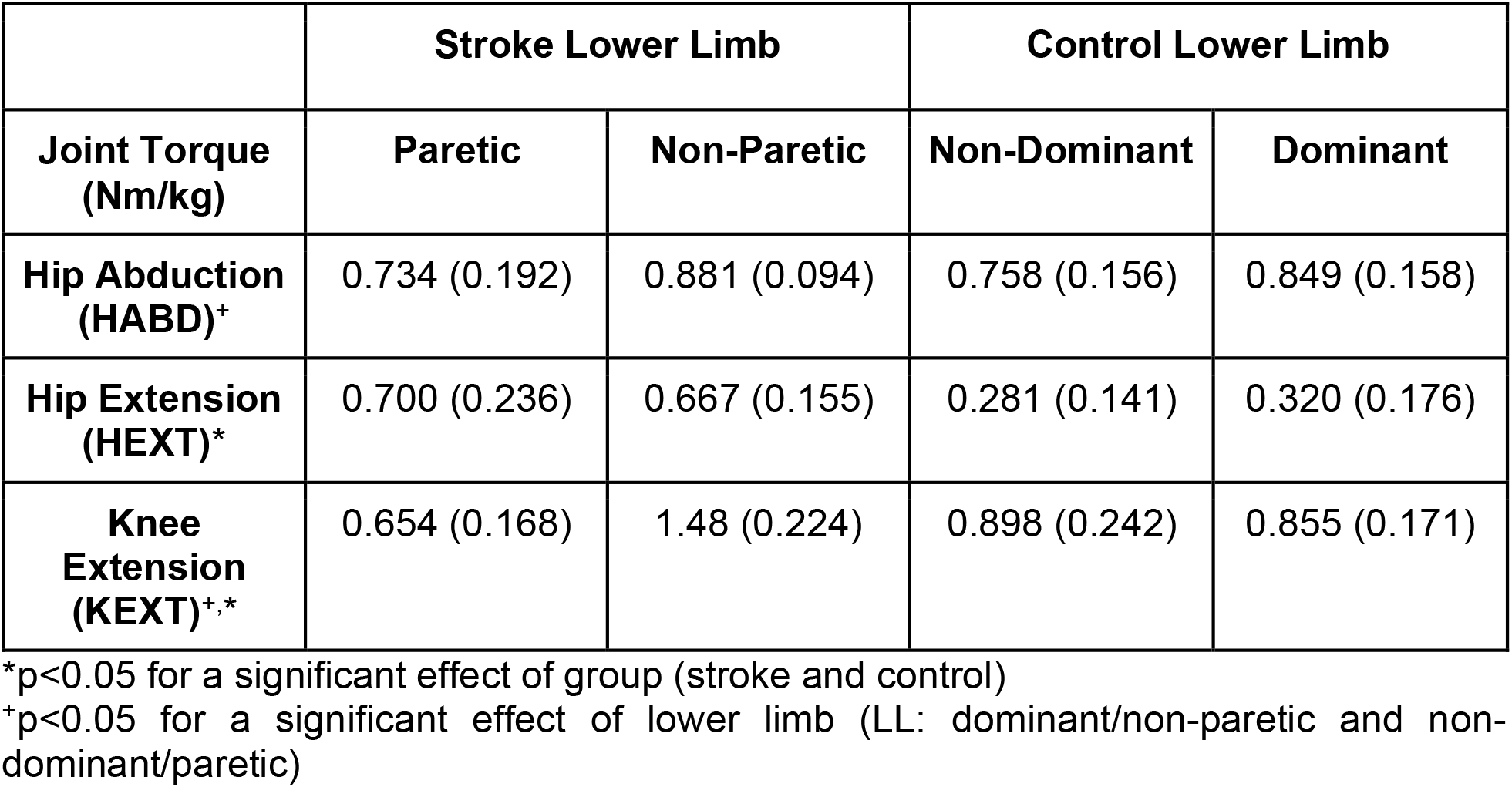
Mean (SD) peak joint torques in hip abduction, hip extension, and knee extension for each leading lower limb during the pull-up phase of step-up.

For the regression model investigating whether peak HEXT and KEXT torque could predict peak HABD torque, results for the stroke paretic limb indicated that only KEXT (b = −0.592, p < 0.001) and the interaction between KEXT and HEXT (b = −0.635, p = 0.002) were significant predictors of HABD. For the stroke non-paretic limb, only KEXT was a significant predictor of HABD (b = 0.179, p = 0.030). There were no significant predictors for the control non-dominant limb, and the control dominant limb showed HEXT (b = 0.557, p = 0.001) and the KEXT by HEXT interaction (b = 0.466, p = 0.022) as significant predictors of HABD.

### III. Joint Kinematics

Joint angle means and standard deviations are displayed in Table 3 and Figure 2. For HABDa, there was no significant main effect of group or LL. For HFLEX, there was no significant effect of group but there was a significant effect of LL (p < 0.001), where the dominant/non-paretic limbs demonstrated greater peak HFLEX angles than the non-dominant/paretic limbs. The interaction between group and LL was also significant (p < 0.001), with the stroke paretic limb exhibiting significantly less peak HFLEX angles than the stroke non-paretic limb (CI: [−21.9 −2.74]). For KFLEX, there was no significant effect of LL but there was a significant effect of group (p = 0.030), where the control group demonstrated greater peak KFLEX angles than the stroke group. The interaction term was also significant (p < 0.001), with the stroke paretic limb exhibiting significantly less peak KFLEX angles than the stroke non-paretic limb (CI: [−43.5 −21.0]), control dominant limb (CI: [−43.5 −22.2]), and control non-dominant limb (CI: [−42.1 −21.8]).

**Table 3.** Mean (SD) peak joint angles in hip abduction, hip flexion, and knee flexion for each leading lower limb during the swing phase of step-up.

|  | Stroke Lower Limb |  | Control Lower Limb |  |
| --- | --- | --- | --- | --- |
| Joint Angles<br>(° Degrees) | Paretic | Non-Paretic | Non-Dominant | Dominant |
| Hip Abduction<br>(HABDa) | 3.65 (2.86) | 8.83 (5.97) | 6.16 (6.58) | 3.97 (4.34) |
| Hip Flexion<br>(HFLEX)<sup>+</sup> | 45.4 (4.88) | 57.7 (8.86) | 51.3 (13.6) | 53.3 (14.5) |
| Knee Flexion<br>(KFLEX)<sup>*</sup> | 49.5 (10.8) | 81.8 (8.48) | 81.5 (4.63) | 82.4 (5.31) |

<sup>\*</sup> $p < 0.05$ for a significant effect of group (stroke and control)
<sup>+</sup> $p < 0.05$ for a significant effect of lower limb (LL: dominant/non-paretic and non-dominant/paretic)

**Figure 2.**
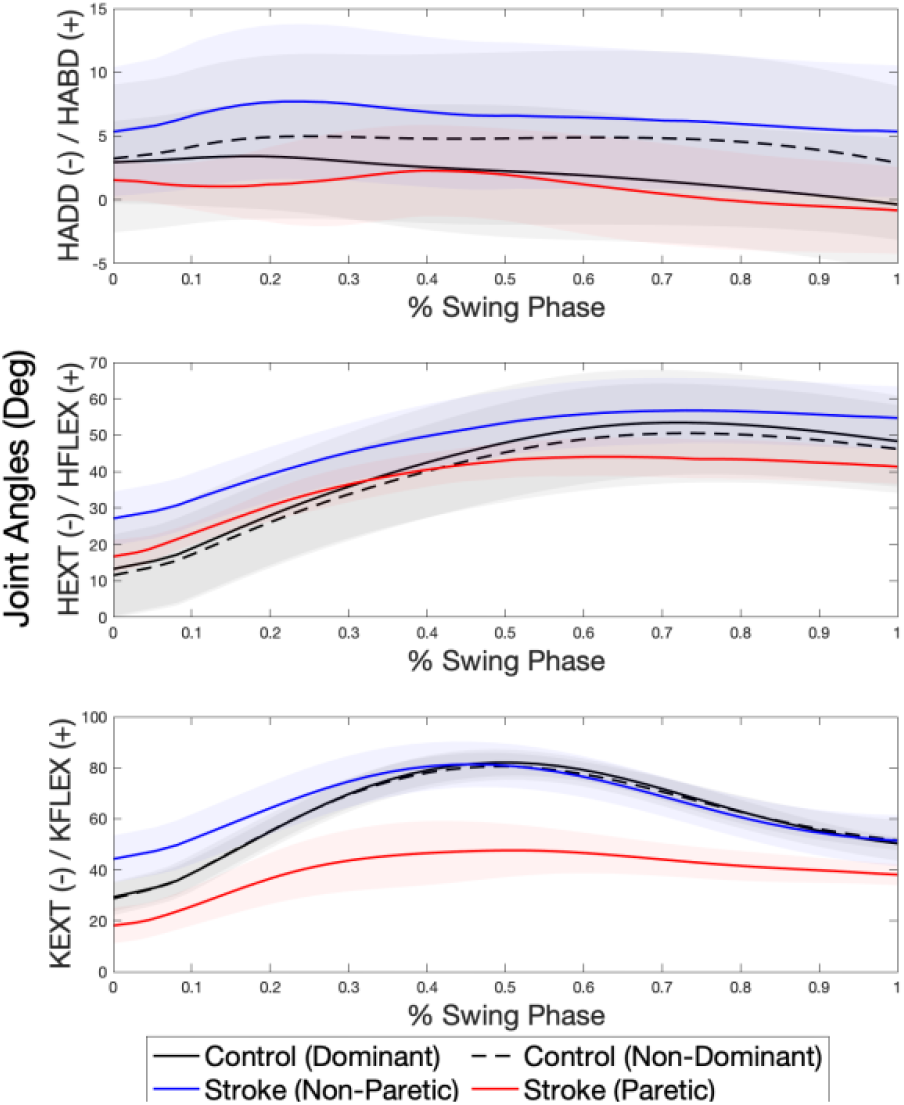
Joint angle traces for the leading leg swinging up onto the step. The x-axis of each subplot is time normalized to 100% of the swing phase and the y-axis is joint angles in degrees. Shaded regions represent one standard deviation. Red line = stroke paretic limb, blue line = stroke non-paretic limb, black dashed line = control non-dominant limb, black solid line = control dominant limb.

## Discussion

This novel study investigated leading hip and knee kinetics and kinematics during the first step of stair ascent in adults with chronic stroke. These two joints are typically the largest contributors to joint angles during swing and joint torque during the pull-up phase of stair ascent, particularly in the frontal and sagittal planes [11-15]. The main finding was that the paretic lower limb exhibited angle differences during swing and torque differences during pull-up stance compared to the non-paretic and control lower limbs. In addition to supporting our hypotheses about task performance, these results reveal insight into the underlying neural consequences of a stroke.

Sánchez et al. quantified hip abduction and knee extension isometric weakness in the paretic limb compared to the non-paretic and control limbs [3]. We confirmed our primary kinetic hypothesis by quantifying significant decreases in these paretic limb joint torques when pulling up onto a step, similar to what has been previously reported [16]. However, high hip extension torque in the paretic limb compared to the control limbs is a new finding. One possible explanation for this mechanically inefficient strategy during pull-up is distal joint weakness at the knee and ankle [3], which is primarily responsible for reduced plantarflexion moments during stance in level-ground walking [22] and could further explain the reduced knee extension torque in this task. High hip and knee extension torque generation in the non-paretic limb during pull-up could also be interpreted as compensation for lack of ankle push-off power in the paretic trailing limb [23]. Collectively, distal joint weakness implies damage to direct corticospinal pathways, as they project more extensively to distal musculature [24,25].

We also confirmed our secondary kinetic hypothesis that increases in peak knee extension torque would predict decreases in peak hip abduction torque only in the paretic limb. Moreover, while peak hip extension torque was not a significant main effect in the paretic limb, the interaction between peak hip and knee extension torque did significantly predict decreases in peak hip abduction torque. The magnitude of the interaction term coefficient was higher than that of knee extension torque. This finding lends evidence for abnormal torque coupling between the hip adductors and sagittal plane extensors in the paretic limb during stepping, a coupling that was not identified in the non-paretic or control limbs. Abnormal extensor coupling has previously been quantified in isometric studies [2,4] and its presence has been attributed to intrinsic neural structures over joint weakness [2], particularly because of the greater co-activation of hip adductor muscles in the paretic limb compared to the control limbs during hip extension efforts [4]. Feline hindlimb models have connected extensor patterns to activation of bulbospinal structures [26,27], which have been postulated to play a role in intralimb stereotypical coupling following stroke [2,4].

The leading paretic limb had reduced hip and knee flexion angles when swinging onto the step, confirming our kinematic hypothesis. Similarities in hip abduction angles could be due to the fact that these joint angles are relatively small compared to sagittal plane flexion angles. Additionally, hip hiking in the paretic limb is common to achieve proper foot clearance in level-ground gait [28], which may be exaggerated when stepping up. Knee joint stiffness in the paretic limb is often responsible for reduced knee flexion during swing and can be attributed to decreases in amplitude and frequency of biceps femoris and gastrocnemius muscles [29] and overactive muscle reflexes of the rectus femoris muscle [30]. Damaged corticospinal pathways from stroke can give rise to this altered neuromuscular drive, particularly for muscles that serve distal joints.

In summary, we quantified the differences in leading hip and knee biomechanical strategies between the paretic and non-paretic/control limbs, defining the nature of asymmetrical motor impairments from hemiparetic stroke. Altered strategies for step ascent included reduced sagittal plane flexion angles during swing, reduced hip abduction and knee extension torque combined with increased hip extension torque during pull-up stance, and abnormal torque coupling between the hip adductors and sagittal plane extensors. These differences can be linked to the neural consequences of a hemiparetic stroke, including corticospinal damage and upregulation of bulbospinal motor pathways as compensation. Study limitations include low sample size, inclusion of individuals with ischemic strokes only, and variability in chronicity of stroke. This analysis was a section of a larger study; further analysis of the trailing leg may illuminate the impact of the paretic leg in the push-up phase of stance. Overall, our findings can inform interventions for individuals with chronic stroke in navigating uneven terrain to maximize daily community activity. Targeted strategies to ameliorate functionally active range of motion at the knee and to combine hip abduction and sagittal plane extensor moments could improve toe clearance and maximize stance stability on stairs.

## Acknowledgements

The authors would like to sincerely thank Emily Baker, Katharine Coombes, Danielle Fredricks, Linsey Daluga, and Sarah Hilu for their contributions to this project, and Keith Gordon for facilitating the use of the Human Agility Lab. This research did not receive any specific grant from funding agencies in the public, commercial, or not-for-profit sectors.

